# Impact of missing data correlated with labels to be predicted in neurodegeneration classification tasks

**DOI:** 10.1101/2025.01.23.634117

**Authors:** Mithilesh Prakash, Jussi Tohka, Alzheimer’s Disease Neuroimaging Initiative

**Author notes:** **Corresponding Author:** Mithilesh Prakash, Ph.D. Biomedical Image Analysis Group University of Eastern Finland Kuopio Campus, Bioteknia Neulaniementie 2, P.O. Box 1627, 70211 Kuopio, Finland **Email:**. A part of the Data used in the preparation of this article were obtained from the Alzheimer’s Disease Neuroimaging Initiative (ADNI) database (adni.loni.usc.edu). As such, the investigators within the ADNI contributed to the design and implementation of ADNI and/or provided data but did not participate in the analysis or writing of this report. A complete listing of ADNI investigators can be found at: http://adni.loni.usc.edu/wp-content/uploads/how_to_apply/ADNI_Acknowledgement_List.pdf.

## Abstract

We introduce a new subtype of ‘Missing Not at Random’ (MNAR) data, where the missingness is correlated with the labels (*y*) to be predicted, termed *(y)-dependent MNAR*. We demonstrate that this subtype can significantly bias the estimation of performance metrics in typical machine learning tasks. Unbiased error estimation is crucial in predictive modeling to accurately assess model performance, identify potential biases, and ensure generalizability to new, unseen data.

We explore the effects of imputing this new subtype of MNAR and compare it with general missing types, namely Missing at Random (MAR) and Missing Completely at Random (MCAR). Our comparison analysis employs both synthetic and clinical datasets, including the Alzheimer’s Disease Neuroimaging Initiative (ADNI) dataset, the Parkinson’s Progression Markers Initiative (PPMI) dataset, and the Anti-Amyloid Treatment in Asymptomatic Alzheimer’s Disease (A4) dataset. After introducing missingness into the datasets, we trained different classifiers paired with various imputation methods and measured repeated cross-validation test metrics.

Our findings reveal that datasets with non-ignorable missing types (MNAR) exhibit a strong bias compared to ignorable types (MAR and MCAR) in downstream analysis. Non-linear classifiers tend to exploit patterns from imputed data, particularly when the imputed values correlate with the target label (*y*), which can lead to unreliable estimation of the generalization error. Mean and median imputations proved to be more robust than tree-based or gradient boosting methods.

## 1 Introduction

Data is often referred to as the new oil of the current age, and statistical predictive models developed using increasingly larger datasets are fueling the AI revolution in the mid-2020s [1]. The performance of these models depends not only on the quantity of the data but also on its quality [2]. The presence of missing values in a dataset can affect the quality of the data available to the model, which in turn is reflected in the performance metrics of models based on such incomplete datasets. Unbiased error estimation is crucial in predictive modeling to assess a model’s performance accurately [3]. It helps identify potential biases and ensures that the model generalizes well to new, unseen data, ultimately ensuring its real-world applicability. The crux of this work is to quantify potential biases in error estimation when developing predictive models using real-world data that often comes with missing values.

Missingness generally occurs in three types: 1) MCAR (Missing Completely at Random), where the probability of missing data, denoted as *P* (miss.), is unrelated to all variables, both observed and unobserved, such that *P* (miss.|complete data) = *P* (miss.); 2) MAR (Missing at Random), where the probability of missing data is related only to the observed data, meaning *P* (miss.|complete data) = *P* (miss.|obs.); and 3) MNAR (Missing Not at Random), where the probability of missing data is related to the missing variables, implying *P* (miss.|complete data) ≠ *P* (miss.|obs.) [4, 5]. Little and Rubin’s introduction of these types of missingness can be grouped as ignorable (MCAR and MAR) and non-ignorable (MNAR) categories when handling missingness in datasets. However, these traditional definitions of missingness do not separate the distinct roles of dependent (*X*) and independent variables (*y*), shown in Table 1, which are important in machine learning experiments. The focused variant of MNAR follows the general definition of MNAR with dependence on the *X*_missing_ component, while the *y*-dependent and diffused variants have a dependence on the *y* component of the dataset.

**Table 1:** Differentiation of missing data mechanisms based on dependence. For a complete dataset comprising *X*_observed_, *X*_missing_, and *y*, this table illustrates how different mechanisms are influenced by various components. The dataset is divided into dependent *X* and independent variables *y* in the machine learning context.

| | | $X$ | | $y$ |
| --- | --- | --- | --- | --- |
| | | $X_{\text{observed}}$ | $X_{\text{missing}}$ | |
| **MCAR** |  | – | – | – |
| **MAR** |  | ✓ | – | – |
| **MNAR** | *Focused* | – | ✓ | – |
|  | *y-dependent* | – | – | ✓ |
|  | *Diffused* | – | ✓ | ✓ |

Simple methods to overcome missing data, such as dropping incomplete observations (complete case analysis) from analysis can lead to data loss and introduce bias in predictions depending on the sample size [6]. Imputation techniques have been developed to address this incompleteness, often with assumptions about the nature of the missingness in the dataset (Table 2). The primary assumption with imputation techniques is that the missingness is not dependent on the missing values and can be resolved using the existing data. However, in machine learning experiments as we will demonstrate in this work, to properly estimate the generalization error an assumption of *y*-independence is additionally needed. Moreover, employing auto-imputation abilities integrated into model training methods [7], such as tree-based classifiers, increases the risk of violating these primary assumptions, leading to models that may not generalize consistently and potentially misleading the performance estimators. The situation is further complicated because there is no established method to detect or confirm a missingness type based on the data to missingness pattern [8]. Moreover, the datasets used to train machine learning models are typically collected through separate sampling (see [9]) so that missingness mechanism can rarely be assumed to be truly informative of underlying phenomenon of biomedical interest.

**Table 2:**
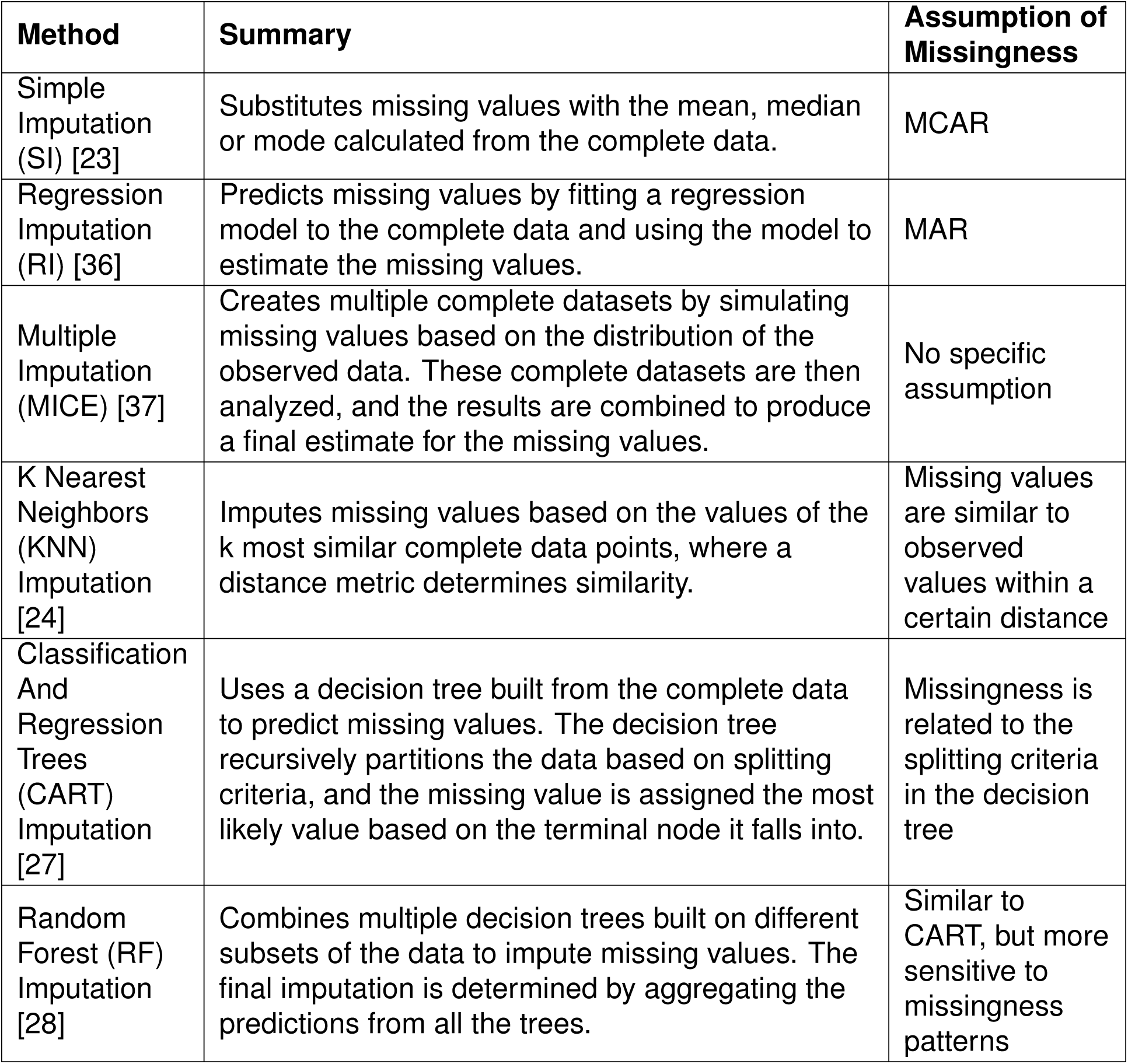
Summary of Imputation algorithms.

| Method | Summary | Assumption of Missingness |
| --- | --- | --- |
| Simple Imputation (SI) [23] | Substitutes missing values with the mean, median or mode calculated from the complete data. | MCAR |
| Regression Imputation (RI) [36] | Predicts missing values by fitting a regression model to the complete data and using the model to estimate the missing values. | MAR |
| Multiple Imputation (MICE) [37] | Creates multiple complete datasets by simulating missing values based on the distribution of the observed data. These complete datasets are then analyzed, and the results are combined to produce a final estimate for the missing values. | No specific assumption |
| K Nearest Neighbors (KNN) Imputation [24] | Imputes missing values based on the values of the k most similar complete data points, where a distance metric determines similarity. | Missing values are similar to observed values within a certain distance |
| Classification And Regression Trees (CART) Imputation [27] | Uses a decision tree built from the complete data to predict missing values. The decision tree recursively partitions the data based on splitting criteria, and the missing value is assigned the most likely value based on the terminal node it falls into. | Missingness is related to the splitting criteria in the decision tree |
| Random Forest (RF) Imputation [28] | Combines multiple decision trees built on different subsets of the data to impute missing values. The final imputation is determined by aggregating the predictions from all the trees. | Similar to CART, but more sensitive to missingness patterns |

Understanding the mechanism of missingness is paramount when modeling health-care data due to the potential severity of the consequences in clinical or medical applications [10]. This understanding becomes particularly important when dealing with multimodal or multicenter datasets, such as those used to model diseases in neurodegeneration studies, where more than one type of missingness can occur [11, 12]. For instance, in a study on Alzheimer’s disease, missing data can occur in multimodal data (e.g., MRI, PET scans, genetic data, cognitive tests) collected from multiple centers. The missingness could be MCAR (e.g., due to random equipment failure), MAR (e.g., older participants might be less likely to complete certain tests), or MNAR (e.g., participants with severe symptoms might be more likely to drop out of the study or be excluded).

The issue of missingness in neurodegeneration datasets has been extensively studied using popular public datasets such as the Alzheimer’s Disease Neuroimaging Initiative (ADNI). Campos et al. [13] demonstrated that imputed datasets with MCAR missingness outperform complete case analysis on ADNI, but did not consider MAR and MNAR. Building on this, Garcia et al. [14] showed that MAR is more prone to bias than MCAR when modeling longitudinal data, but did not study MNAR. Zhou et al. [15] conducted MNAR experiments using the ADNI dataset, addressing the less studied MNAR missingness. Later, McCombe et al. [16] conducted a comprehensive simulation experiment on ADNI considering both MAR and MCAR. In a more recent study, Chandrasekaran et al. [17] utilized MICE to enhance performance on imputing datasets with MAR missingness in longitudinal ADNI data, but did not consider MNAR or MCAR.

Despite these studies, a significant limitation is the reliance on curated clinical datasets, where the true performance of the disease prediction model remains uncertain. In such cases, having a simulated dataset with known performance metrics can be a significant asset. A classification task pairs well with a random dataset where the association of noise (predictors) to random labels can be controlled with a Bayes rate. This allows us to set a theoretical limit for a perfect classifier, and data leakages due to imputation or violation of assumptions can be observed through deviations from the norm.

The definitions introduced by Gomer et al. expand the MNAR subtypes to include situations where missingness depends only on the missing values, termed focused MNAR, and where missingness depends on both the missing values and the observed data, termed diffuse MNAR [18]. We build on this definition to recognize a situation where missingness depends on the target label, which we call *y*-dependent MNAR. We also demonstrate that this new variant is distinct from other types and variants of MNAR, and highlight the importance of identifying it in the modeling process. We want to emphasize that, if the distinction between predictors and labels is absent, this new variant is just MAR and does not pose a problem. However, if the distinction is made, as in modeling in machine learning, the missingness type changes to MNAR because the missing values depend on the values themselves, which can be problematic.

In this study, we aim to demonstrate how different missing mechanisms impact down-stream analysis. We expand the MNAR definition, introduce missingness in the data, and then pair the missing data with popular imputation methods. Finally, we model the imputed data with classifiers, all on a performance-capped dataset. Next, we showcase an actual clinical dataset where our updated MNAR definition was first observed in the Anti-Amyloid Treatment in Asymptomatic Alzheimer’s Disease (A4) dataset. A performance boost on imputed data was observed when distinguishing amyloid positivity in the study population. Finally, we experiment with public and private neurodegenerative datasets and analyze the performance.

The primary objective of this study is to increase awareness within the research community and explore possible variants in missingness definitions. Additionally, with the automation in data analysis, researchers may inadvertently use auto-imputation methods and find it difficult to understand the failure of a model’s generalizability.

## 2 Methods

We set up a binary classification problem where **X** = [*X*_1_*, X*_2_*, …, X_k_*] represents a vector of predictor variables, and *y* ∈ {0, 1} is the binary target variable. The relationship can be expressed as:

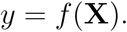

The goal is to learn the function *f* that maps **X** to *y* based on training data 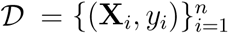. We introduced various kinds of missingness into the data and trained classifiers with this imputed data to understand how different missingness mechanisms caused bias in the estimation of the classifier’s generalization error and how to overcome it.

We briefly introduce the datasets (Table 3), the missingness mechanisms (Figure 1 and Table 4), as well as the imputation and classification steps used for the downstream analysis.

**Figure 1:**
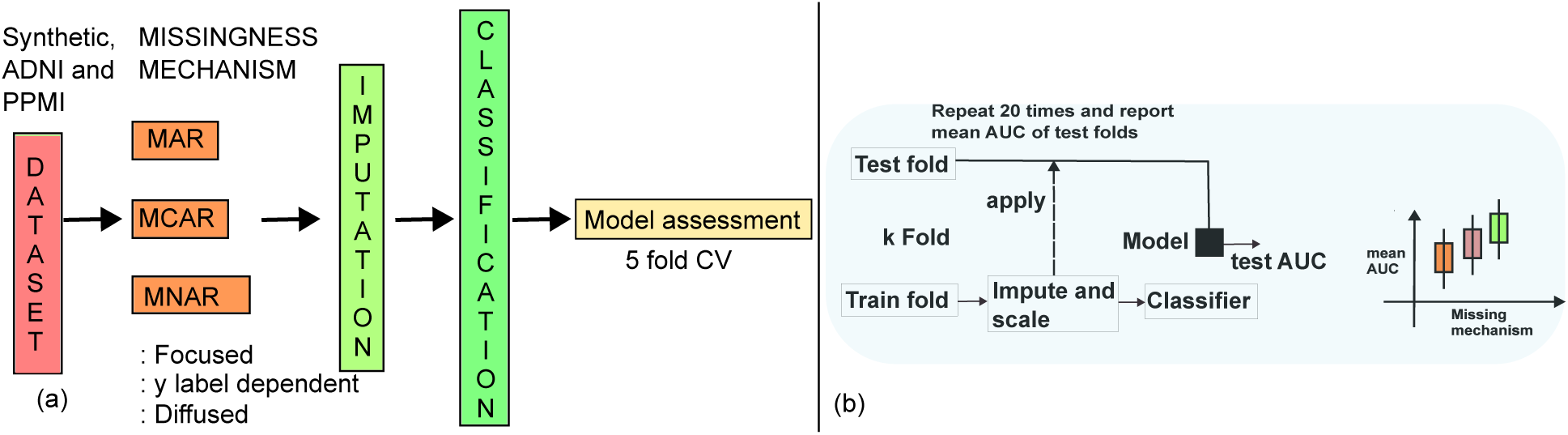
Study outline (a) and modeling overview (b).

**Table 3:** Summary of datasets with their characteristics (*X*) and targets (*y*)

| Dataset | N | Sex (M/F) | Age<br>(mean±sd)<br>yrs | Number of<br>features | Binary target ($y$) |
| --- | --- | --- | --- | --- | --- |
| Synthetic | 1000 | - | - | 50 | Random with Bayes rate<br>at 0.50, 50% each. |
| ADNI | 1691 | 814/877 | 72 ± 7 | 6 | Healthy or mild cognitive<br>impairment (MCI):<br>(779/912) |
| PPMI | 1453 | 900/553 | 62 ± 10 | 10 | Healthy controls or<br>Parkinson's disease (PD):<br>(267/1186) |
| A4 | 4486 | 2663/1823 | 71 ± 4 | 73 | Amyloid eligibility status<br>(positive/negative):<br>1323/3163 |

**Table 4:** Table of Missingness Types and Proportions.

| Type of Missingness | Dataset name | Proportion % |
| --- | --- | --- |
| Missing completely at Random | mcar20 | 20 |
|  | mcar40 | 40 |
|  | mcar80 | 80 |
| Missing at Random | mar20 | 20 |
|  | mar40 | 40 |
|  | mar80 | 80 |
| Missing not at Random | *y-dependent* |  |
|  | mnarY20 | 20 |
|  | mnarY40 | 40 |
|  | mnarY80 | 80 |
|  | *Focused* |  |
|  | mnarF20 | 20 |
|  | mnarF40 | 40 |
|  | mnarF80 | 80 |
|  | *Diffused* |  |
|  | mnarD20 | 20 |
|  | mnarD40 | 40 |
|  | mnarD80 | 80 |

### 2.1 Datasets

The synthetic and clinical datasets employed in this study are summarized in Table 3.

#### 2.1.1 Synthetic data

The synthetic dataset used in this study consists of 1000 samples and 50 features. The features are generated from a standard normal distribution, ensuring that each feature follows a Gaussian distribution with a mean of 0 and a standard deviation of 1. The target variable, *y*, is a binary variable generated using a binomial distribution with a success probability of 0.50. This means that each sample has an equal probability of being assigned a class label of 0 or 1 and the Bayes rate for the classification task is 0.50.

#### 2.1.2 ADNI data

A part of the data used in the preparation of this article was obtained from the Alzheimer’s Disease Neuroimaging Initiative (ADNI) database (http://adni.loni.usc.edu) [19]. Launched in 2003 as a public-private partnership led by Principal Investigator Michael W. Weiner, MD, ADNI aims to test whether serial magnetic resonance imaging (MRI), positron emission tomography (PET), other biological markers, and clinical and neuropsychological assessments can be combined to measure the progression of mild cognitive impairment (MCI) and early Alzheimer’s disease (AD). For up-to-date information, see http://www.adni-info.org.

With ADNI data, including participants with cognitively normal (CN) or with mild cognitive impairment (MCI) status, the task was to decide whether a participant is categorized as CN (*y_i_* = 0) or suffering from MCI (*y_i_* = 1) at the baseline visit. The dataset used is derived from the ADNI MERGE table. The selected features include age, gender, years of education, APOE4 genotype, intracranial volume (ICV), and hippocampal volume. Rows with missing values were excluded to ensure data completeness because we simulated the missingness of different types in our experiments. Binary variables were created for gender and cognitive status, resulting in a dataset where the features are age, years of education, APOE4 genotype, intracranial volume, hippocampal volume, and binary gender, while the target variable is the binary cognitive status. The RIDs of the participants are provided in the supplement.

#### 2.1.3 PPMI data

A part of the data used in the preparation of this article was obtained on [2024-01-29] from the PPMI database (http://www.ppmi-info.org/access-dataspecimens/download-data, *RRID:SCR 006431*) [20]. For up-to-date information on the study, visit http://www.ppmi-info.org. We use only the data from the baseline visit. The selected predictor features are gender, years of education, handedness, age, Montreal Cognitive Assessment (MoCA) score, Benton Judgment of Line Orientation Test (BJLOT) score, Hopkins Verbal Learning Test (HVLT) immediate recall, delayed recall, recognition scores, body mass index (BMI). The target variable is the participant classification (Healthy vs Parkinson’s disease). Rows with missing values were excluded to ensure data completeness because we will simulate the missingness of different types in our experiments.

#### 2.1.4 A4 data

A part of the data used in the preparation of this article was obtained from the Anti-Amyloid Treatment in Asymptomatic Alzheimer’s (A4) database (https://www.a4studydata.org/). The A4 screening dataset represents the largest cohort of cognitively unimpaired older individuals with amyloid imaging and comprehensive cognitive testing available to date. For detailed information, see (https://www.clinicaltrials.gov/study/NCT02008357).

The A4 study includes volumetric MRI data combined with demographic information, cognitive scores, APOE genotype, and amyloid eligibility (our target variable) [21]. The selected MRI features include various volumetric measurements, along with the Hippocampal Occupancy (HOC) Score. HOC is defined as the ratio of hippocampal volume to the sum of hippocampal and inferior lateral ventricle volumes. The demographic features include age, years of education, retirement status, marital status, and gender. Cognitive scores included PACC, Digit Symbol, FCSRT (2xFree + Cued), Logical Memory Delay, and MMSE. Rows with missing values in the MRI data were identified, and binary variables were created for gender, retirement status, marital status, and APOE genotype. This dataset has missing values correlated with our target variable and hence we use this to showcase our new variant of MNAR.

### 2.2 Experiments

#### 2.2.1 Missingness mechanism

We explored five mechanisms of missing data: *MCAR* (Missing Completely at Random), *MAR* (Missing at Random), and three types of *MNAR* (Missing Not at Random) (Figure 1). Each mechanism introduced missingness in different ways, impacting the analysis and interpretation of the data. For the *i*-th individual, let *M_i_*∈ {0, 1}*^k^* denote the missingness indicator vector and we write **x***_i_* = [**x**_(_*_obs_*_)_*_i_,* **x**_(_*_miss_*_)_*_i_*], where **x**_(_*_obs_*_)_*_i_* denotes the values of the predictors which are observed and **x**_(_*_miss_*_)_*_i_* the values which are missing. Note that in our supervised learning scenario, we always assume that the target variables *y_i_* are observed.

*MCAR* occurs when the probability of missing data on a variable is independent of any other observed or unobserved data. Mathematically, this can be expressed as:

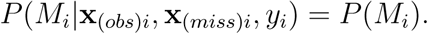

In this scenario, data is missing in completely random fashion, and any analysis remains unbiased. In our implementation, missing values are introduced randomly across the dataset based on a specified probability, affecting only a certain percentage of observations.

*MAR* is characterized by the probability of missing data on a variable being related to some of the observed predictor data but not the missing data itself. This can be formulated as:

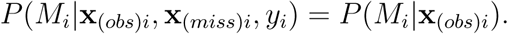

Here, the missingness depends on the observed data **x**_(_*_obs_*_)_*_i_*, but not on the missing data **x**_(_*_miss_*_)_*_i_*. For instance, if the missingness in one variable is dependent on the values of another observed variable, the data is considered MAR. In our implementation, missing values are introduced based on a threshold in one column and applied to another column, simulating the dependency on observed data. Only a certain percentage of observations that meet the threshold criteria are randomly affected.

*MNAR* occurs when the probability of missing data on a variable is related to the value of the variable itself, even if the value is missing or it is the target variable and *P* (*M_i_*|**x**_(_*_obs_*_)_*_i_,* **x**_(_*_miss_*_)_*_i_, y_i_*) cannot be simplified in the general case. We further categorize MNAR into three types based on different conditions:

- *Focused MNAR*: Missingness is introduced based on a threshold in the data, focusing on specific values within the dataset. Missing values are introduced where the data exceeds a certain threshold, with only a certain percentage of these observations being randomly affected. Missingness does not depend on the target variable, i.e., *P* (*M_i_*|**x**_(_*_obs_*_)_*_i_,* **x**_(_*_miss_*_)_*_i_, y_i_*) = *P* (*M_i_*|**x**_(_*_miss_*_)_*_i_*).
- *y-dependent MNAR*: Missingness is introduced solely based on the value of the target variable *y*: *P* (*M_i_*|**x**_(_*_obs_*_)_*_i_,* **x**_(_*_miss_*_)_*_i_, y_i_*) = *P* (*M_i_*|*y_i_*). Here, missing values are introduced in the data where the target variable equals 1. Again, only a certain percentage of these observations are randomly affected.
- *Diffused MNAR*: Missingness is introduced based on a combination of a threshold in the data and the value of a target variable *y*: *P* (*M_i_*|**x**_(_*_obs_*_)_*_i_,* **x**_(_*_miss_*_)_*_i_, y_i_*) = *P* (*M_i_*|**x**_(_*_miss_*_)_*_i_, y_i_*) Specifically, missing values are introduced where the data exceeds a certain threshold and the target variable equals 1. Only a certain percentage of these qualified observations are randomly affected.

The proportion of randomly affected observations was controlled by a parameter, allowing for flexibility in the degree of missingness introduced (Table 4). We implemented the above definitions in a Python class. Additionally, we utilized the `pyampte` package to introduce missingness similar to our definitions for comparisons with the Synthetic Dataset [22].

#### 2.2.2 Imputation and Learning Algorithms

The imputation methods employed included not imputing (specifically for the GB classifier), mean, median [23], KNN (2 nearest neighbors to prevent overfitting) [24, 25], MICE (Multiple Imputation by Chained Equations) [26], and a tree-based method [27, 28].

We studied several classifiers to solve the formulated classification tasks. These learning algorithms included the Gradient Boosting Classifier (GB), which builds an ensemble of decision trees, the Linear Discriminant Analysis (LDA), a linear classifier that projects data onto a lower-dimensional space, the Calibrated Support Vector Classifier (SVC) with a linear kernel and regularization parameter *C* = 0.1, and Random Forest Classifier (RF). RF, an ensemble method using multiple decision trees, was employed with a maximum depth of 2 because this setting enhances computational efficiency and prevents overfitting, ensuring better generalization on unseen data [29].

We used the implementations available in the `scikit-learn` package for imputation methods * and learning algorithms ^†^. The hyperparameters were in default settings unless specified above.

#### 2.2.3 Generalization Error Estimation

We evaluated the classifiers using repeated (n=20) k-fold cross-validation (k=5, *stratified* for equal proportions of target labels in all folds). The mean AUC values, aggregated from all repetitions and cross-validation test folds, were used as the performance measures. It is important to note that while each cross-validation run had non-overlapping test folds, the same samples could appear in different test folds across repetitions.

We emphasize that it is important to perform the splitting of the data into training and test folds prior to imputation in order to avoid data leakage. However, we also performed an evaluation without a data split prior to imputation. This was done to investigate the effect of performing imputation on the entire data set before dividing it into training and testing sets. This allowed for a comparison of the performance of the classifiers when trained on imputed data with and without a prior data split, providing an understanding of the bias in error estimation by this (incorrect) modeling.

The baseline performance of the ADNI and PPMI datasets was established using a gradient boosting classifier with default parameter settings on complete datasets, i.e., without any missing data introduced. For the A4 dataset, we marked the baseline from the published result from Petersen et al. [30].

The code implementing the reported experiments is available at: https://github.com/mithp/MNAR_experiments_2024.git

## 3 Results

We discuss the main findings of the experiments under its own header.

### 3.1 Types of Missingness in the Datasets Had a Major Effect on Downstream Analysis

The type of missingness was observed to affect downstream analysis. MNAR showed a positive performance bias (synthetic dataset: observed AUC range of 0.50 − 1.00 where AUC 0.50 would be correct), proving data leakage compared to MAR and MCAR (synthetic dataset: observed AUC range of 0.47 − 0.53) (Figure 2). This performance difference can be attributed to the realignment of predictors after imputation (Figure 3). These effects were observed in both synthetic (Figure 4) and clinical datasets (Figures 5 and 6), though they were less pronounced in the latter due to inherent feature correlations.

**Figure 2:**
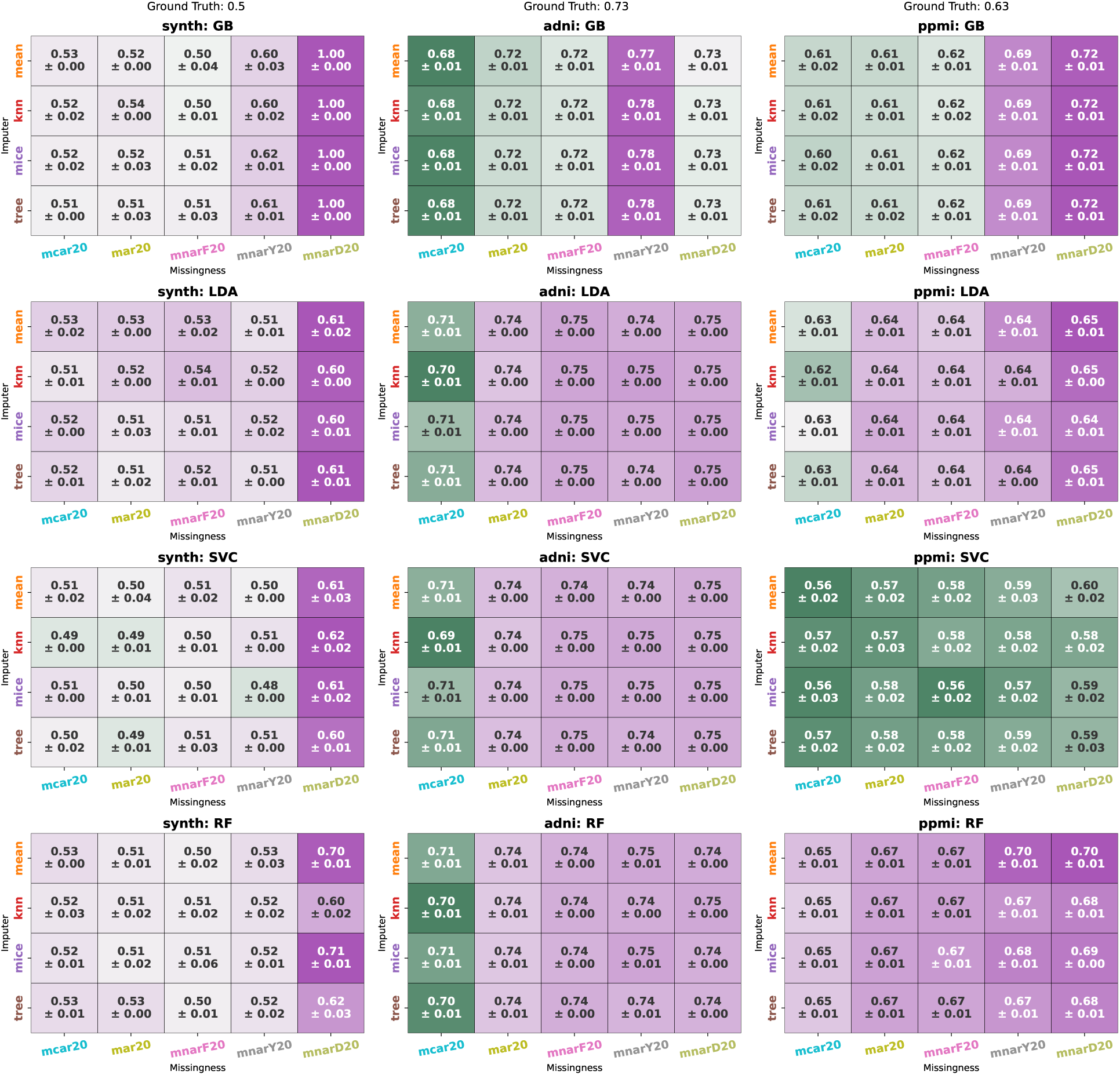
Summary of Experiments: This heatmap shows the mean test set Area Under the Curve (AUC) (±) standard deviation of the values for each dataset (column-wise). The Missing at Random (MAR, 20% missing data: `mar20`) and Missing Completely at Random (MCAR, 20% missing data: `mcar20`) scenarios are not affected, with AUC close to ground truth for each dataset. In contrast, the Missing Not at Random scenarios (focused: `mnarF20`, y-label dependent: `mnarY20`, and diffused: `mnarD`, both at 20% missing data) show deviations from ground truth AUC values. Each column displays classifiers in its own subplots, with each row representing the imputation method used alongside the classifiers. The classifiers used are Gradient Boosting (GB), Linear Discriminant Analysis (LDA), Support Vector Classifier (SVC), and Random Forest (RF). The imputation methods include mean, Multiple Imputation by Chained Equations (MICE), k-Nearest Neighbors (kNN), and tree-based methods.

**Figure 3:**
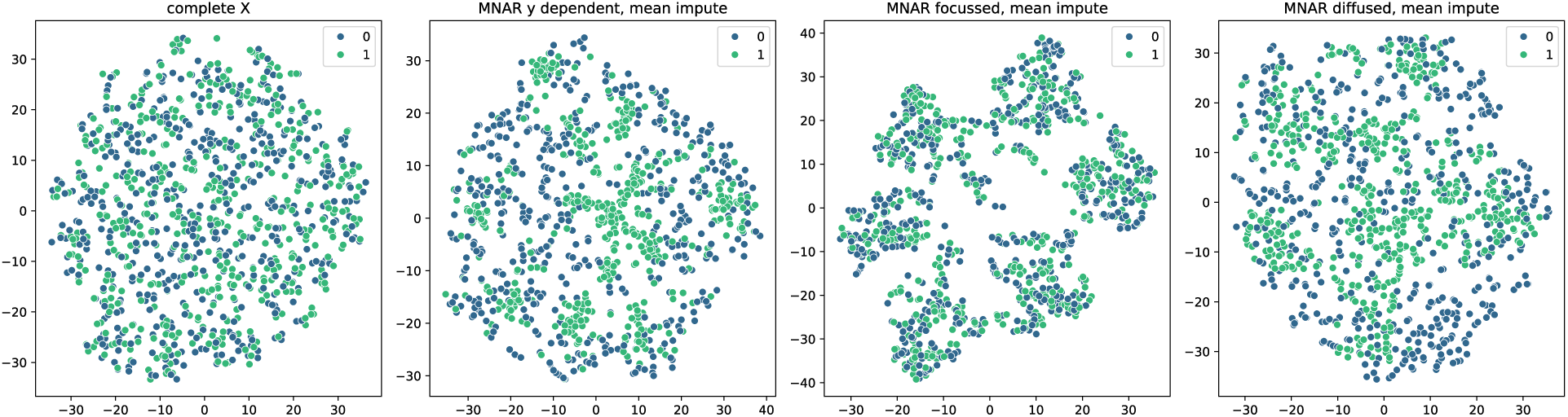
Missingness Types with Mean Imputation in Synthetic Dataset: t-distributed Stochastic Neighbor Embedding (t-SNE) Visualization: The rearrangement of (*y*) labels (0/1) is different for various missingness patterns of Missing Not at Random (MNAR) types, changing the distribution in the (*X*) space and thus influencing the downstream classifier performance. This visualization uses mean imputation to handle missing data (set at 80%).

**Figure 4:**
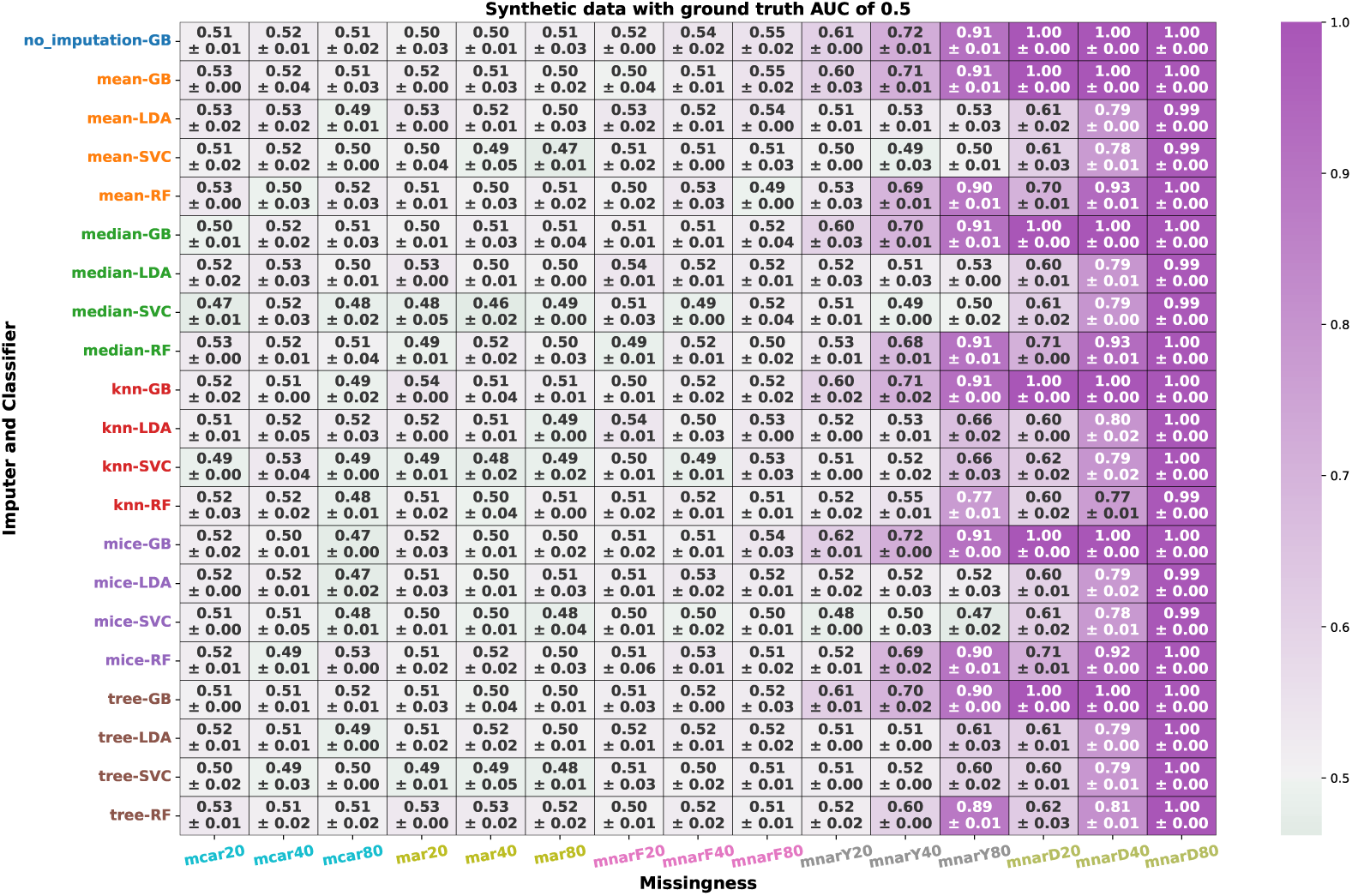
**Experiments on Synthetic Data** were conducted with the Bayes rate set at 0.50 (ground truth). AUC values (mean test set (±) standard deviation) above 0.50 indicate potential data leakage and false gains. Missingness mechanisms include Missing Not at Random (MNAR) scenarios: focused (`mnarF20`, `mnarF40`, `mnarF80`), y-label dependent (`mnarY20`, `mnarY40`, `mnarY80`), and diffused (`mnarD20`, `mnarD40`, `mnarD80`); Missing at Random (MAR) (`mar20`, `mar40`, `mar80`); and Missing Completely at Random (MCAR) (`mcar20`, `mcar40`, `mcar80`). The postfixes 20, 40, and 80 indicate 20%, 40%, and 80% missing data, respectively. AUC of test folds are grouped by missingness type.

**Figure 5:**
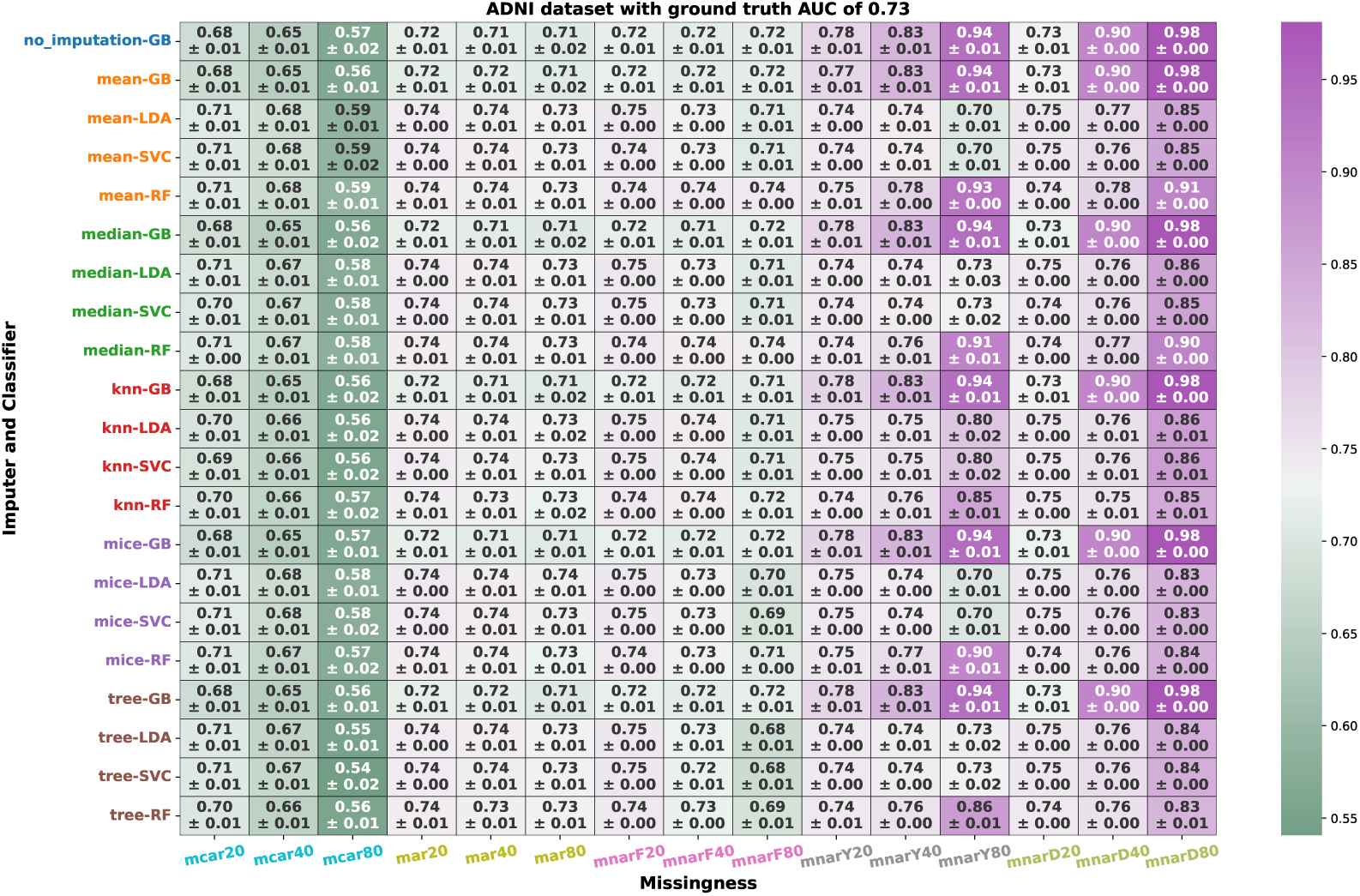
**Experiments on ADNI data** were conducted with the ground truth AUC of 0.73 in predicting the control vs. mild cognitive impairment. See Fig. 4 for other notation.

**Figure 6:**
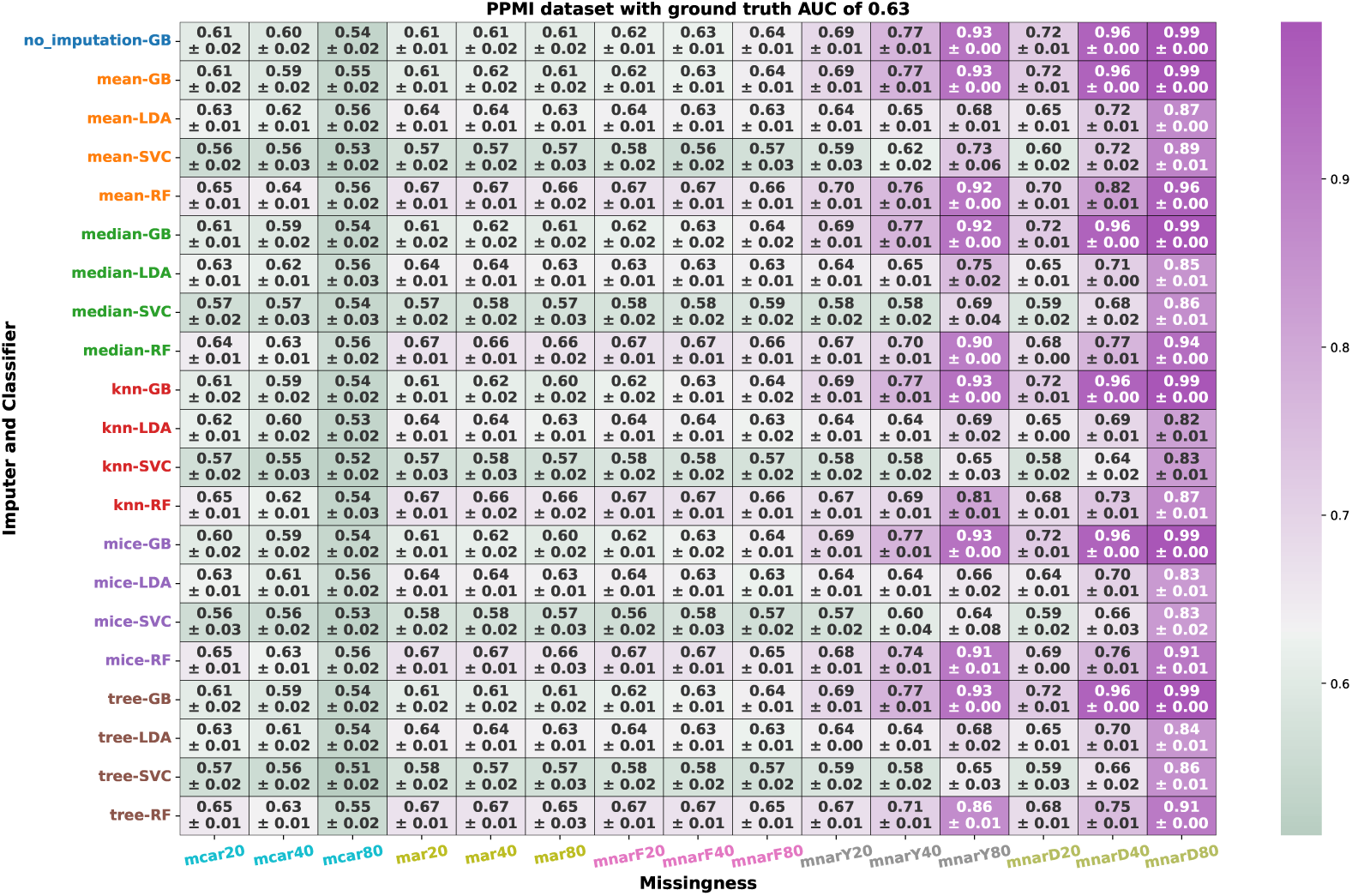
**Experiments on PPMI data** were conducted with the ground truth AUC of 0.63 in predicting the control vs. mild Parkinson’s Disease. See Fig. 4 for other notation.

### 3.2 Choosing a classifier revealed that linear classifiers are more robust to imputed data than tree-based classifiers

Linear classifiers displayed robustness (synthetic dataset, mean imputation, `mnarY40`: AUC range of 0.49 − 0.53) to different missingness mechanisms compared to tree-based classifiers (synthetic dataset, mean imputation, `mnarY40`: AUC range of 0.60 − 0.73). Linear classifiers were not sensitive to the *y*-dependent missingness type of MNAR, whereas tree-based classifiers readily picked up the missingness pattern (lighter vs. darker shades in Figure 4). Similarly, in A4 dataset (MNAR: *y*-dependent missingness pattern, Figure 7) where GB and RF (AUC range of 0.81−0.97) picked up imputation patterns that matched to to AB+ group labels, whereas LDA and SVC were more robust (AUC range of 0.49 − 0.82) to artefactual missingness pattern (Figure 7, ground truth for A4 dataset was 0.72).

**Figure 7:**
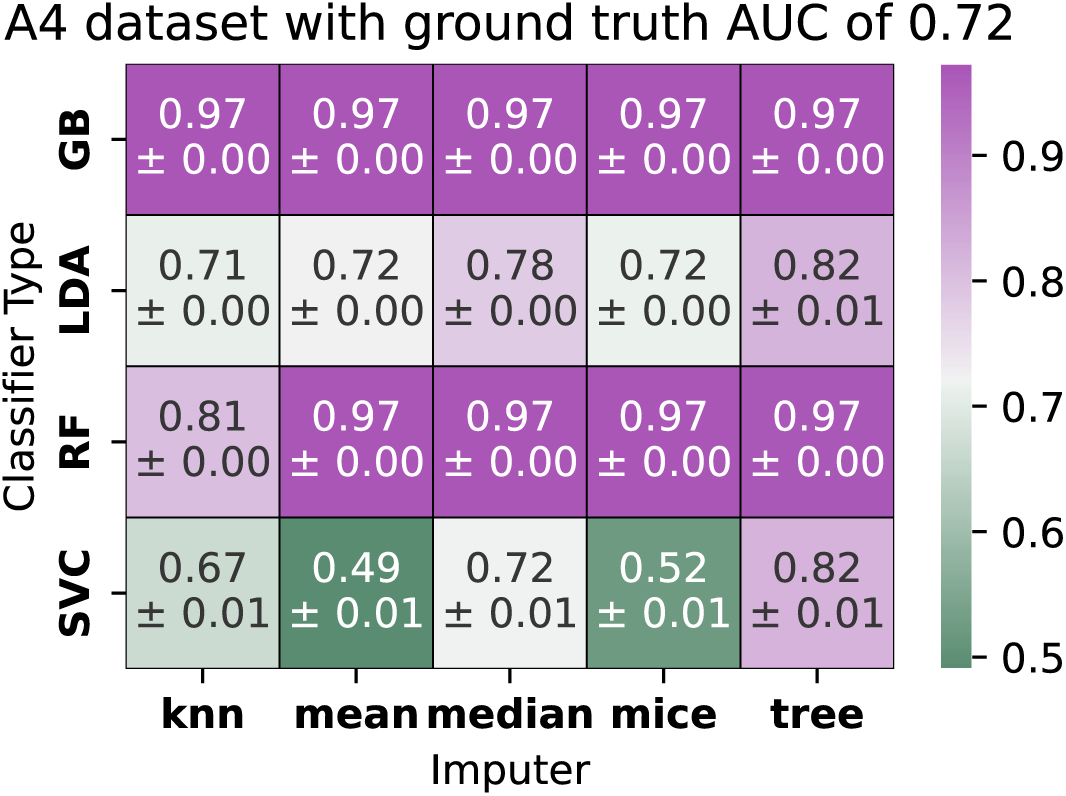
**The A4 dataset** where missingness is ignored, imputed, and data modeled to exhibit the bias in Missing Not at Random (MNAR) type of missingness in a clinical dataset. The heatmap shows mean test set (±) standard deviation.. An AUC of 0.72 is used as a ground truth from Petersen et al. [30]. See Fig. 4 for other notation.

### 3.3 The choice of imputation methods had a minor effect on the downstream analysis

The impact of imputation methods on classifier performance was overshadowed by the effects of missingness types and classifier choice. However, mean and median imputation methods were less sensitive when combined with linear classifiers compared to other imputation methods and classifier combinations.

The imputation methods showed the least variation when the missingness mechanisms were MAR, MCAR, and MNAR-focused for both synthetic and clinical datasets, and this behavior extended to the MNAR-focused group as well. For the y-dependent MNAR type at 40% and 80% missing data (`mnarY40` and `mnarY80`), KNN imputation showed robustness when paired with linear classifiers as observed with synthetic data (linear (0.52 − 0.66) vs. non-linear(0.55 − 0.91), Figure 4) and ADNI data (linear (0.75 − 0.80) vs. non-linear(0.76 − 0.94), Figure 5), while only impacting RF in the PPMI dataset. In the A4 dataset, mean and MICE imputation showed larger variations in the AUC with the baseline for the SVC classifier compared to median and KNN imputation. However, these variations were minimized for the LDA classifier. Overall, mean, median, and MICE were consistent across the experiments.

### 3.4 Missing percentages have different effects depending on the types of missingness and classifier combinations

The increase in missing value percentage affected the random data (synthetic data) and clinical data (ADNI and PPMI) experiments differently. In the former dataset, the MAR, MCAR, and MNAR focused variants were unaffected, while in the latter, performance metrics decreased as anticipated for the MAR and MCAR types (AUCs in ADNI: ∼0.75 to ∼0.60; PPMI: ∼0.65 to ∼0.50). However, MNAR variations showed an increase in performance (AUCs in ADNI: ∼0.75 to ∼0.95; PPMI: ∼0.70 to ∼0.95).

*y*-dependent MNAR and diffused MNAR affected both dataset types similarly. The effect was less pronounced in clinical datasets (Δ ∼ 0.25) than in synthetic datasets (Δ ∼ 0.45) due to feature correlations.

### 3.5 Splitting (vs. not) the data into train and test before imputation had the least effect on synthetic datasets and showed some minor effects on clinical datasets

Not splitting the data before imputation did not result in drastic changes (Δ in AUCs), but linear classifiers (LDA and SVC) showed false gains when data splitting was ignored (Figures 4, 5, and 6 vs. Supplementary Figures **??**, **??**, and **??**) when paired with mean, median, and KNN imputation methods.

## 4 Discussion

We studied supervised learning and generalization error estimation in the case of missing data. We experimented with three subtypes of MNAR (`mnarF`, `mnarY`, `mnarD`) and demonstrated data leakage due to imputation across all experiments, which manifests as false gains (estimated performance above the expected/true performance with complete data). In contrast, MAR and MCAR maintained expected performance estimates, with no clear bias.

We demonstrated the real-world relevance of this work using the A4 clinical dataset, which has missing data due to selective data sampling. These studies on performance estimation in the case of missing data are especially important not only because there is no gold standard method to identify the missingness mechanism [31], but also due to the high potential for unintentional bias caused by data leakage from imputed datasets.

Our findings can be summarized as follows: 1)The types of missingness in the datasets had a major effect on downstream analysis, as observed by the clear bias in performance metrics for MNAR variants; 2) It was found that linear classifiers are more robust to imputed data than tree-based classifiers; 3) The choice of imputation methods had a minor effect on the downstream analysis; 4) Missing percentages have different effects depending on the types of missingness and classifier combinations; 5) Splitting the data into train and test before imputation had the least effect on synthetic datasets and showed some minor effects on clinical datasets.

Studies simulating missingness patterns and their effects on downstream analysis cannot be reliably tested by employing only research datasets. This is because complete datasets are rare, and the baseline or benchmark cannot be reliably established. New progress in modeling techniques can advance the baseline, as witnessed in the year-on-year accuracy improvements in the computer vision MNIST (Modified National Institute of Standards and Technology) challenge [32]. As a partial answer to this challenge, we relied on a synthetic dataset to reliably establish a theoretical limit that can be set on a perfect classifier.

It is accepted that when there is missingness in the dataset, the usual loss in predictive power results due to data loss [33]. However, a problematic trend can occur when the missingness is of a non-ignorable variant and imputation is performed, potentially leading to issues such as data leakage. The MNAR experiments demonstrate this when compared to other ignorable types of missingness (MAR and MCAR) in the dataset. Amongst the MNAR subtypes, we see that when the missing values depend on the missing value itself (`mnarF`), the effect is similar to ignorable missingness. This is because, in both correlated datasets (ADNI and PPMI) and random datasets (Synthetic Data), the prediction power is either compensated through imputation if correlated, or it deteriorates due to actual data loss. Additionally, the imputation clearly does not change the predictors. This kind of MNAR (`mnarF`) could be reliably imputed as witnessed by the results. However, the behavior is different for y-dependent (`mnarY`) and diffused (`mnarD`) versions of MNAR because there is an underlying correlation with the prediction label, and this can be seen due to the rearrangement of data into clusters which could be picked up in downstream analysis. We demonstrated this behavior with the A4 dataset, and we attribute the missingness pattern observed in this dataset to be y-label dependent (`mnarY`). In such scenarios, we recommend a simple correlation test between the missingness pattern as a mask (0/1) and the target label to help determine the nature of the pattern, although no standard statistical test exists.

The choice of classifier is important when considering imputed datasets because the pattern in the dataset changes. For example, with mean imputation, if the missingness pattern are correlated with the label to be predicted, then non-linear classifiers tend to exploit these patterns. This can lead to unreliable estimation of the generalization error by cross-validation based error estimators. The decision boundary from linear classifiers clearly did not exploit the imputed values or the new pattern, but the non-linear classifiers readily picked up the pattern due to their inherent characteristics. The underlying decision trees in RF and the boosting of weak learners to strong learners in GB could explain the improved pattern recognition [34], compared to the decision boundaries of their linear counterparts.

We observe that these linear vs. non-linear properties also affected the imputation effects. Mean and median replacements for missing values were more robust than tree-based or auto imputation by gradient boosting. Together with MICE, mean and median imputations are recommended considering their robustness to overall effects.

In our view, the variations in the percentages of missing values amplified the effects of the missingness type, classifier, and imputation combination. As the missing percentages increase, the underlying patterns become stronger and easier for non-linear classifiers to detect. Conversely, for linear combinations, the data loss for ignorable missing types results in the expected loss of predictive power.

One important step to prevent data leakage is to avoid double-dipping by splitting the training and testing folds and applying the information gained from the training data to transform the testing data [35]. For example, this applies to scaling or imputing. We wanted to see if this made a difference in the experiments, but the missingness overwhelmed the effect of data leakage due to ignoring the data split when imputing the dataset. Nonetheless, data splitting should be exercised as part of best practices to reliably generalize the model to independent or unseen data.

To conclude, our experiments revealed that the type of missingness profoundly impacts downstream analysis. Non-linear classifiers tend to exploit patterns from imputed data, particularly when the imputed values correlate with the target label, which can be problematic because it may lead to unreliable estimation of the generalization error. Mean and median imputations proved to be more robust than tree-based or GB methods.

## 5 Acknowledgements

This study was funded by the Research Council of Finland grants (349184, 346934, 358944 (Flagship of Advanced Mathematics for Sensing Imaging and Modeling)), grant 351849 from the Research Council of Finland under the frame of ERA PerMed (“Pattern-Cog”) and Saastamoinen Foundation grants 2023.

Data collection and sharing for this project was funded by the Alzheimer’s Disease Neuroimaging Initiative (ADNI) (National Institutes of Health Grant U01 AG024904) and DOD ADNI (Department of Defense award number W81XWH-12-2-0012). ADNI is funded by the National Institute on Aging, the National Institute of Biomedical Imaging and Bio-engineering, and through generous contributions from the following: AbbVie, Alzheimer’s Association; Alzheimer’s Drug Discovery Foundation; Araclon Biotech; BioClinica, Inc.; Biogen; Bristol-Myers Squibb Company; CereSpir, Inc.; Cogstate; Eisai Inc.; Elan Pharmaceuticals, Inc.; Eli Lilly and Company; EuroImmun; F. Hoffmann-La Roche Ltd and its affiliated company Genentech, Inc.; Fujirebio; GE Healthcare; IXICO Ltd.; Janssen Alzheimer Immunotherapy Research & Development, LLC.; Johnson & Johnson Pharma-ceutical Research & Development LLC.; Lumosity; Lundbeck; Merck & Co., Inc.; Meso Scale Diagnostics, LLC.; NeuroRx Research; Neurotrack Technologies; Novartis Phar-maceuticals Corporation; Pfizer Inc.; Piramal Imaging; Servier; Takeda Pharmaceutical Company; and Transition Therapeutics. The Canadian Institutes of Health Research is providing funds to support ADNI clinical sites in Canada. Private sector contributions are facilitated by the Foundation for the National Institutes of Health (http://www.fnih.org). The grantee organization is the Northern California Institute for Research and Education, and the study is coordinated by the Alzheimer’s Therapeutic Research Institute at the University of Southern California. ADNI data are disseminated by the Laboratory for Neuro Imaging at the University of Southern California.

PPMI — a public-private partnership — is funded by the Michael J. Fox Foundation for Parkinson’s Research and funding partners, including 4D Pharma, Abbvie, AcureX, Al-lergan, Amathus Therapeutics, Aligning Science Across Parkinson’s, AskBio, Avid Radio-pharmaceuticals, BIAL, BioArctic, Biogen, Biohaven, BioLegend, BlueRock Therapeutics, Bristol-Myers Squibb, Calico Labs, Capsida Biotherapeutics, Celgene, Cerevel Therapeutics, Coave Therapeutics, DaCapo Brainscience, Denali, Edmond J. Safra Foundation, Eli Lilly, Gain Therapeutics, GE HealthCare, Genentech, GSK, Golub Capital, Handl Therapeutics, Insitro, Janssen Neuroscience, Jazz Pharmaceuticals, Lundbeck, Merck, Meso Scale Discovery, Mission Therapeutics, Neurocrine Biosciences, Neuropore, Pfizer, Piramal, Prevail Therapeutics, Roche, Sanofi, Servier, Sun Pharma Advanced Research Company, Takeda, Teva, UCB, Vanqua Bio, Verily, Voyager Therapeutics, the Weston Family Foundation, and Yumanity Therapeutics.

The A4 Study was a secondary prevention trial in preclinical Alzheimer’s disease, aiming to slow cognitive decline associated with brain amyloid accumulation in clinically normal older individuals. The A4 Study was funded by a public-private-philanthropic partnership, including funding from the National Institutes of Health-National Institute on Aging, Eli Lilly and Company, Alzheimer’s Association, Accelerating Medicines Partner-ship, GHR Foundation, an anonymous foundation, and additional private donors, with in-kind support from Avid Radiopharmaceuticals, Cogstate, Albert Einstein College of Medicine, and the Foundation for Neurologic Diseases. The companion observational Longitudinal Evaluation of Amyloid Risk and Neurodegeneration (LEARN) Study was funded by the Alzheimer’s Association and GHR Foundation. The A4 and LEARN Studies were led by Dr. Reisa Sperling at Brigham and Women’s Hospital, Harvard Medical School, and Dr. Paul Aisen at the Alzheimer’s Therapeutic Research Institute (ATRI) at the University of Southern California. The A4 and LEARN Studies were coordinated by ATRI at the University of Southern California, and the data are made available under the auspices of the Alzheimer’s Clinical Trial Consortium through the Global Research & Imaging Platform (GRIP). The complete A4 Study Team list is available at: https://www.actcinfo.org/a4-study-team-lists/. We would like to acknowledge the dedication of the study participants and their study partners who made the A4 and LEARN Studies possible..

The computational analysis was run on the servers provided by Bioinformatics Center https://bioinformatics.uef.fi/biowhat/BIC/bic_101/, University of Eastern Finland, Finland. During the preparation of this work, the authors used Microsoft Copilot from Internet Explorer to improve language and readability. After using this tool, the authors reviewed and edited the content as needed. The authors take full responsibility for the content of the publication.

## 6 Author contributions

MP and JT conceptualized and designed the study. MP performed the analysis. Both authors contributed to the writing and approved the final manuscript.

## Footnotes

* `SimpleImputer(mean), SimpleImputer(median), KNNImputer(), IterativeImputer(ExtraTreesRegressor)`

† `HistGradientBoostingClassifier(), LinearDiscriminantAnalysis(), CalibratedClassifierCV(SVC()), RandomForestClassifier()`

